# Kinetic Monitoring Of Neuronal Stress Response To Proteostasis Dysfunction

**DOI:** 10.1101/2021.05.24.445437

**Authors:** Angel J. Santiago-Lopez, Ken Berglund, Robert E. Gross, Claire-Anne N. Gutekunst

## Abstract

Proteostasis dysfunction and activation of the unfolded protein response (UPR) are characteristic of all major neurodegenerative diseases. Nevertheless, although the UPR and proteostasis dysfunction has been studied in great detail in model organisms like yeast and mammalian cell lines, it has not yet been examined in neurons. In this study, we applied a viral vector-mediated expression of a reporter protein based on a UPR transcription factor, ATF4, and time-lapse fluorescent microscopy to elucidate how mouse primary neurons respond to pharmacological and genetic perturbations to neuronal proteostasis. In *in vitro* models of endoplasmic reticulum (ER) stress and proteasome inhibition, we used the ATF4 reporter to reveal the time course of the neuronal stress response relative to neurite degeneration and asynchronous cell death. We showed how potential neurodegenerative disease co-factors, ER stress and mutant α-synuclein overexpression, impacted neuronal stress response and overall cellular health. This work therefore introduces a viral vector-based reporter that yields a quantifiable readout suitable for non-cell destructive kinetic monitoring of proteostasis dysfunction in neurons by harnessing ATF4 signaling as part of the UPR activation.

## INTRODUCTION

The protein homeostasis (proteostasis) network comprises the processes by which cells control protein production, maintenance, and degradation to support cellular function (Hipp et al., 2019). To maintain proteostasis, the endoplasmic reticulum (ER) serves as an intracellular sensor that captures signals associated with proteostasis dysregulation and induces corrective measures through a series of signaling events collectively known as the unfolded protein response (UPR) (Hetz and Saxena, 2017). The UPR in the ER can be activated by ER stress – a biological consequence of the imbalance between proper protein handling (i.e., folding, trafficking, degrading) and the cellular ER capacity. The UPR is broadly defined as part of the integrated stress response (ISR), which expands to stressors beyond the ER, such as amino acid deficiency, viral infection, and heme deprivation (Pakos-Zebrucka et al., 2016).

Proteostasis dysfunction is one of the leading theories to describe cell-autonomous drivers of neurodegeneration in human neurodegenerative diseases. Stress in the ER and subsequent activation of the UPR has been associated with the presence of misfolded protein aggregates in neurodegenerative diseases (Abisambra et al., 2013; Hoozemans et al., 2012; Nijholt et al., 2012) and neuroprotective interventions have been designed by directly modulating UPR signaling *in vivo* (Bugallo et al., 2020; Mercado et al., 2018; Radford et al., 2015). Thus, understanding of neuronal responses to impairment in proteostasis such as the UPR remains a central area of study in neuroscience with potential impact on drug development.

In mammalian systems, cellular signaling in the UPR is governed by three signaling programs mediated by the activating transcription factor-6 (ATF6), and the kinases IRE1α, and PERK. As a downstream effector of the UPR, the activating transcription factor-4 (ATF4), also known as the cAMP-response element binding protein 2 (CREB-2), is selectively translated during cellular stress following the phosphorylation of eIF2α by the kinases of the ISR (PERK, GCN2, HRI, or PKR) (Moon et al., 2018; Vattem and Wek, 2004). ATF4 activates a transcriptional program aimed at combating oxidative stress, restoring amino acid metabolism, and, in conditions of unresolved stress, executing apoptosis. As such, the preferential translation of ATF4 is regarded as a biological signature arising from the cellular stress response (Wortel et al., 2017).

The biological outcomes of the UPR are commensurate with the level of proteostasis dysfunction and are strongly dependent on the time (or phase) of induction. Importantly, early events in the UPR are generally considered cytoprotective. However, upon unresolved cellular stress, the PERK arm of the UPR, in coordination with IRE1α (Fink et al., 2018; Lin et al., 2007), activates a molecular switch to induce cell death via apoptosis, thereby preventing the presence of cells with unresolved stress to remain in an organism (Frakes and Dillin, 2017; Lin et al., 2009). In this scenario, the CCAAT-enhancer-binding protein homologous protein (CHOP), a downstream product of PERK, triggers cell death as an apoptotic gene (Galehdar et al., 2010). Therefore, understanding the natural progression of the cellular stress response has been instrumental for mapping the timescale of stress signaling activity relative to cellular fate (Walter et al., 2015).

Studies evaluating UPR activation have traditionally employed protein quantification methods for activated UPR kinases or mRNA quantification of UPR target genes such as *ATF3* and *Trib3*. Notably, such conventional biochemical and molecular biology methods have intrinsic limitations in assessing spatial and temporal profiles of cellular stress in live cells as well as limited resolution for discerning between individual cellular responses. The advent of reporter cell lines and luciferase-based stress reporters have enabled a more detailed study of the kinetic aspect of cell fate decisions during conditions of cellular stress (Walter et al., 2015). These tools, however, have not yet been adapted to neuronal systems. Neuronal models such as those established from primary sources better recapitulate neuronal physiology and function as they retain characteristics from their tissue of origin (Gordon et al., 2013). However, primary neurons are generally challenging to transfect in sufficient numbers for high-throughput evaluation via fluorescence microscopy and are also vulnerable to assays requiring multistep cell culture manipulations. These limitations have collectively hindered progress to directly monitor and quantify the kinetics of stress responses in live neurons.

In this study, we sought to visualize stress responses in live neuronal cultures by implementing a viral vector-based reporter construct based on the 5’ untranslated region (UTR) of the ATF4 UPR effector gene. By harnessing this selective translational event, we conducted long-term live-cell imaging studies to examine the chronic stress responses evoked by pharmacological induction of ER stress and proteasome inactivation. Building on these results, we then probed whether the overexpression of A53T mutant α-synuclein, a mutant protein linked to early-onset Parkinson’s Disease, elicited a neuronal stress response, or influenced neuronal susceptibility to ER stress. Collectively, our data demonstrates the feasibility of an ATF4-based cellular assay to conduct kinetic studies of cellular stress following proteostasis dysfunction in neurons.

## METHODS

### Cell culture

Human embryonic kidney (HEK) 293T cells were maintained in a humidified 5% CO_2_ atmosphere at 37 °C in complete medium consisting of Dulbecco’s Modified Eagle’s medium (DMEM) supplemented with 10% fetal bovine serum, 2 mM L-glutamine, 100 international units (IU)/mL penicillin, and 100 µg/mL streptomycin (pen-strep).

### Primary neuronal cultures

Flat bottom black 96 well plates (Greiner CELLSTAR®) were coated with fresh 100 µg/mL poly-L-lysine (Sigma) aqueous solution and incubated at 37°C for at least 12 h. Coated wells were rinsed once with Dulbecco’s phosphate buffer saline (DPBS) and allowed to air dry for 2 h prior to use. Primary neurons were isolated from CD-1 mouse embryos on embryonic day 18 in accordance with approved protocols by the Emory University Institutional Animal Care and Use Committee, and National Institutes of Health guidelines. Brain cortices were dissected and dissociated using Worthington’s Biochemical Papain Dissociation System (Worthington Biochemical Corporation) following manufacturer’s instructions and previously published protocols (Santiago-Lopez et al., 2018). Freshly dissociated neurons were plated in plating medium consisting of Neurobasal™ Plus base medium supplemented with 5% fetal bovine serum, 1 µg/mL gentamycin, and 2 mM Glutamax™-I. Complete medium change was performed 12 h post-plating to maintenance medium consisting of Neurobasal™ Plus base medium supplemented with B-27 Plus, 2 mM Glutamax™-I, and 1% pen-strep. Partial medium changes were performed once a week thereafter with maintenance medium. Cortical neurons were considered mature and optimal for experiments at 20-25 days *in vitro* (DIV).

### Plasmid construction and vector production

To generate stress-dependent expression vectors, cDNA comprising the first 384 bp of the human *ATF4* transcript including the 5’UTR and the initial coding region flanked by the *EcoRI* and *AgeI* restriction sites was synthetized using a commercial service (gBlocks, Integrated DNA Technologies) and cloned into an adeno-associated virus (AAV) 2 transfer plasmid with enhanced green fluorescent protein (EGFP) using conventional molecular biology techniques (Gutekunst et al., 2016). A similar construct lacking the ATF4 cassette was produced as a control for the constitutive expression of GFP. Transgene expression in both constructs is driven by the chicken β-actin (CBA) promoter for ubiquitous expression in mammalian cells. The correct sequences were confirmed by restriction enzyme digestion analyses and DNA sequencing. Expression characteristics of each plasmid were evaluated in HEK 293T cells by transient transfection using Lipofectamine™ 3000 (Thermo Fisher Scientific).

AAV vectors of the DJ hybrid serotype were produced by following previously published protocols (Grieger et al., 2006; Gupta et al., 2020). Briefly, adherent HEK 293T cells were expanded and triple transfected with plasmids encoding for viral capsid, helper virus, and transgene of interest. After 3 d, cells were harvested and lysed by sodium deoxycholate followed by repeated freezing-thawing. Viral particles were purified by an iodixanol density gradient followed by buffer exchange and concentration in 100,000 MWCO filter units. Titer was determined by qPCR to be in the range of 10^12^ viral genomes (vg)/mL. For α-synuclein overexpression studies, lentivirus expression vectors encoding human A53T mutant α-synuclein fused to the red fluorescent protein mKate2 driven by the eIF1α promoter were manufactured by VectorBuilder Inc and reported to have a titer of 10^12^ vg/mL.

### Pharmacological treatments

Thapsigargin (Sigma) was solubilized in DMSO and working solutions were prepared in Neurobasal Plus/B-27 maintenance medium. MG132 (Abcam) was handled in a similar fashion. From working solutions, drugs were added directly to the cells at the appropriate final concentration as further detailed in the results section. Solvent-only solutions were used as control for each drug, accordingly. Culture medium was not replaced after treatment to prevent additional disturbances.

### Image acquisition and analysis

Live-cell imaging experiments were conducted using a Cytation 5 Cell Imaging Multi-Mode Reader (Biotek Inc.) equipped with a gas controller for CO_2_ control and 4-zone temperature incubation with condensation control. Fluorescent images were acquired with either a 10X or 20X phase contrast objectives using either the DAPI, GFP, or CY5 built-in filter cubes with LED. The image acquisition routine was established in the Gen5 Plus software to follow a kinetic read taking at least 12 images in the same x- and y-coordinates for each well throughout the course of each time-lapse session.

Raw images were subjected to semi-automated pixel-based analysis using the Gen5 image analysis function. For reporter analysis, threshold settings were adjusted to capture GFP positive neurons. Image analysis metrics (GFP intensity, GFP area, and GFP integrated intensity) were calculated for each image in an experiment using the same threshold and masking settings. For neurite area quantification, phase contrast images were analyzed to quantify cell body area which was then subtracted from the total cell culture area. Calculated metrics were normalized to the corresponding first reading (t_0_) to express a relative rate of change from baseline.

### Immunocytochemistry

Primary neuronal cultures were fixed with 4% paraformaldehyde for 15 min and rinsed with D-PBS. Following permeabilization with 0.1% Triton X-100 for 5 min, cells were blocked with 4% normal serum for 30 min. Cultures were incubated overnight at 4 °C with primary antibodies followed by three rinses in PBS. Processed samples were incubated with appropriate secondary antibodies for 1 h at room temperature, counterstained with Hoechst 33342, and rinsed with PBS. The following antibodies were used: mouse anti-Tau-1 (1:400, Millipore Sigma), rabbit anti-ubiquitin (1:500, Agilent Dako), rabbit anti-β III tubulin (1:500, Abcam), and donkey anti-mouse or goat anti-rabbit Alexa Fluor® Plus 647 (1:1000, ThermoFisher Scientific).

### Lactate dehydrogenase (LDH) measurement

Lactate dehydrogenase (LDH) release was measured by using the CyQUANT LDH Cytotoxicity Assay (ThermoFisher Scientific) on collected cell culture supernatant according to manufacturer’s instructions. Absorbance was measured at 490 nm and 680 nm using the Cytation 5 plate-reader function. Results are reported as 680 nm-corrected absorbance normalized to untreated control cultures.

### Statistics

Statistical analysis was performed in Graph Pad Prism 7. Data sets were tested for normality using the D’Agostino and Pearson test. For population comparisons not following a normal distribution, statistics were calculated using a two-tailed non-parametric Mann-Whitney test. Multiple group comparisons were analyzed by the Kruskal-Wallis test. Results are reported as mean ± SEM.

## RESULTS

### AAVs with ATF4-based translational control allow for stress-dependent transgene expression in neurons

The selective translation of ATF4 is a convergence point of the ISR following proteostasis dysfunction and activation of the UPR. To harness this event for enabling stress-dependent transgene expression in neurons (Figure 1A), we first adapted ATF4 reporter by cloning the regulatory domain of *ATF4* including the 5’UTR and first 28 amino acids upstream of EGFP. Initial characterization of the resulting construct in HEK293 cells revealed stable reporter suppression following insertion of *ATF4* 5’ UTR upstream of EGFP (_ATF4_GFP) compared to constitutively expressed EGFP nder the same promoter (_C_GFP) (Figure S1A, S1B). Consistent with previous reports (Walter et al., 2015), induction of ER stress with 1 µM thapsigargin (TG) in HEK293 cells increased intensity and number of _ATF4_GFP expressing cells (Figure S1C, S1D).

**Figure 1:**
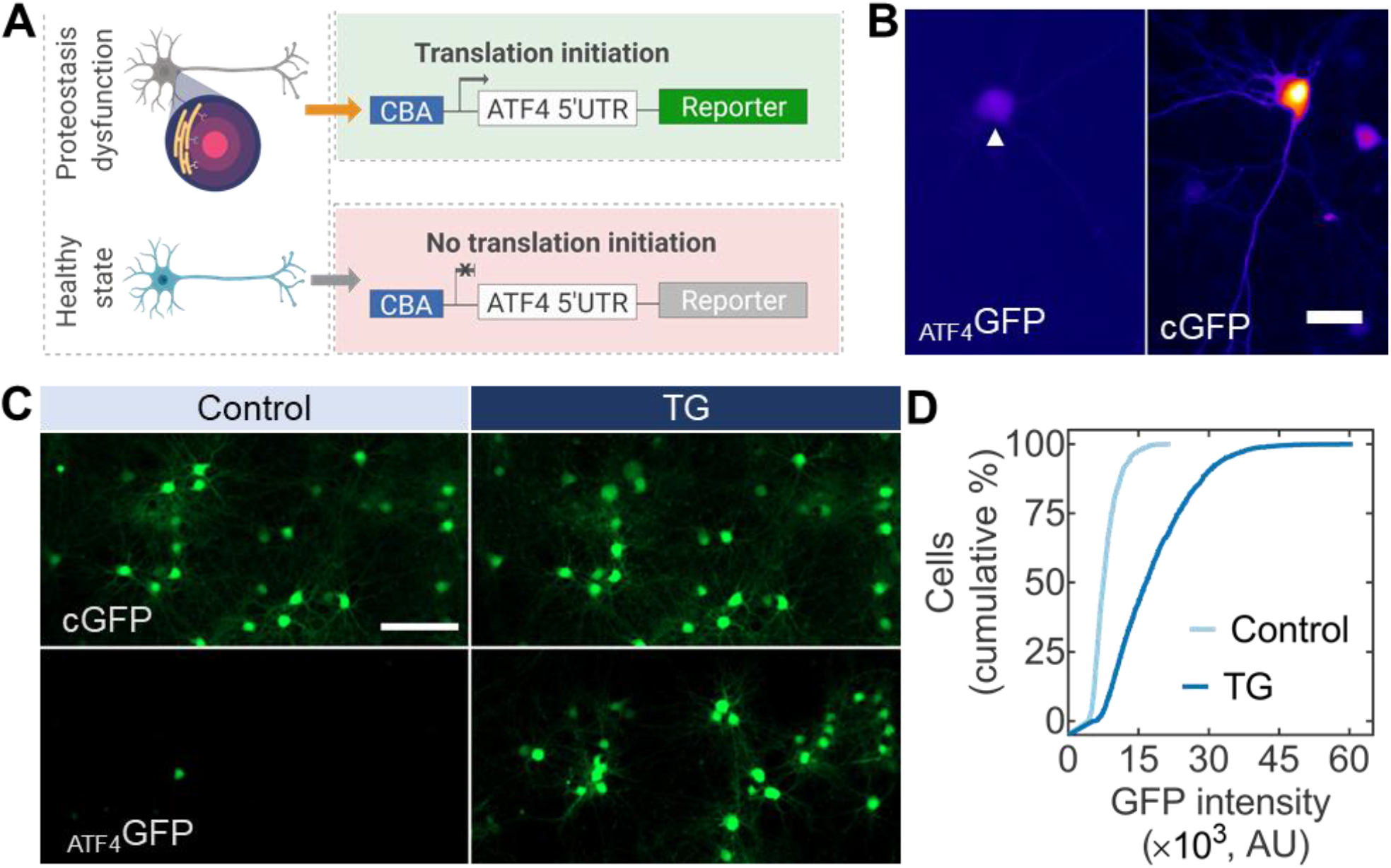
Stress-dependent transgene expression vectors applied in primary neurons. (**A**) AAV vectors with a fluorescent reporter downstream of a translational control operator from the *ATF4* 5’ UTR allow for the assessment of proteostasis dysfunction in neurons. (**B**) Primary cortical neurons under normal culture conditions seven days after viral transduction. Stress-dependent expression (_ATF4_GFP, left, arrow) is substantially lower than constitutively expressed GFP (_C_GFP, right). Scale bar 25 µm. (**C**) Primary neurons expressing _C_GFP (top) or _ATF4_GFP (bottom) untreated (left, control) or treated with TG for 24 h (right, TG). Scale bar 100 µm. (**D**) Population-based quantification of _ATF4_GFP induction from control following TG treatment. n > 3,000 neurons.

Since conventional transfection methods are not compatible for high-throughput assessment of neuronal responses in sufficient numbers (Karra and Dahm, 2010), we opted for AAV vectors pseudotyped with DJ as gene delivery vehicles. The superior *in vitro* transduction performance of the AAV DJ hybrid serotype compared to other AAV variants has been reported previously (Grimm et al., 2008). Similar to HEK293 cells, under normal culture conditions primary cortical neurons transduced with _ATF4_GFP showed a dim transgene expression compared to cultures transduced with _C_GFP (Figure 1B). To evaluate the effectiveness of detecting stress response in neurons, we adopted TG, a classical inducer of ER stress and UPR activation. TG typifies an ER-targeted pharmacological perturbation inhibiting the ER Ca^2+^ ATPase, causing ectopic calcium entry, ER stress, and eventually apoptosis (Treiman et al., 1998). Upon 24 h exposure to 100 nM TG, cumulative distributions of cortical neurons transduced with _ATF4_GFP showed a rightward shift where >50% of the cell population showed at least a two-fold increase in mean _ATF4_GFP intensity compared to control cultures (Figure 1C, 1D). Based on the degree of induction, these results suggest _ATF4_GFP could be used as a metric to capture stress responses in neurons.

### Kinetic monitoring of ER stress in neurons

Compared to other biological systems, the temporal analysis of stress responses arising from proteostasis dysfunction in neuronal systems has not yet been reported. We first opted to characterize the temporal profile of stress response in neurons induced by TG in preparation for evaluating stress responses in more neurobiologically relevant contexts. Even though it is estimated that TG depletes stored Ca^2+^ in a timescale of minutes (Treiman et al., 1998), chronic ER stress has been described to occur ∼6 h post-addition of the ER stressor *in vitro* (Guan et al., 2017). We therefore designed live-cell imaging experiments to cover and exceed the time window for chronic ER stress responses. For the kinetic study of cellular stress in primary neurons, images were acquired every 2-3 h to prevent blue light-induced phototoxicity. Representative images at 0 h (baseline) and 20 h following treatment with TG of cortical neurons show a clear induction of _ATF4_GFP, primarily localized to the cell body (Figure 2A, left panel). On the other hand, neurons transduced with _C_GFP displayed general upregulation of GFP, irrelevant to the treatment (Figure 2A, right panel).

**Figure 2:**
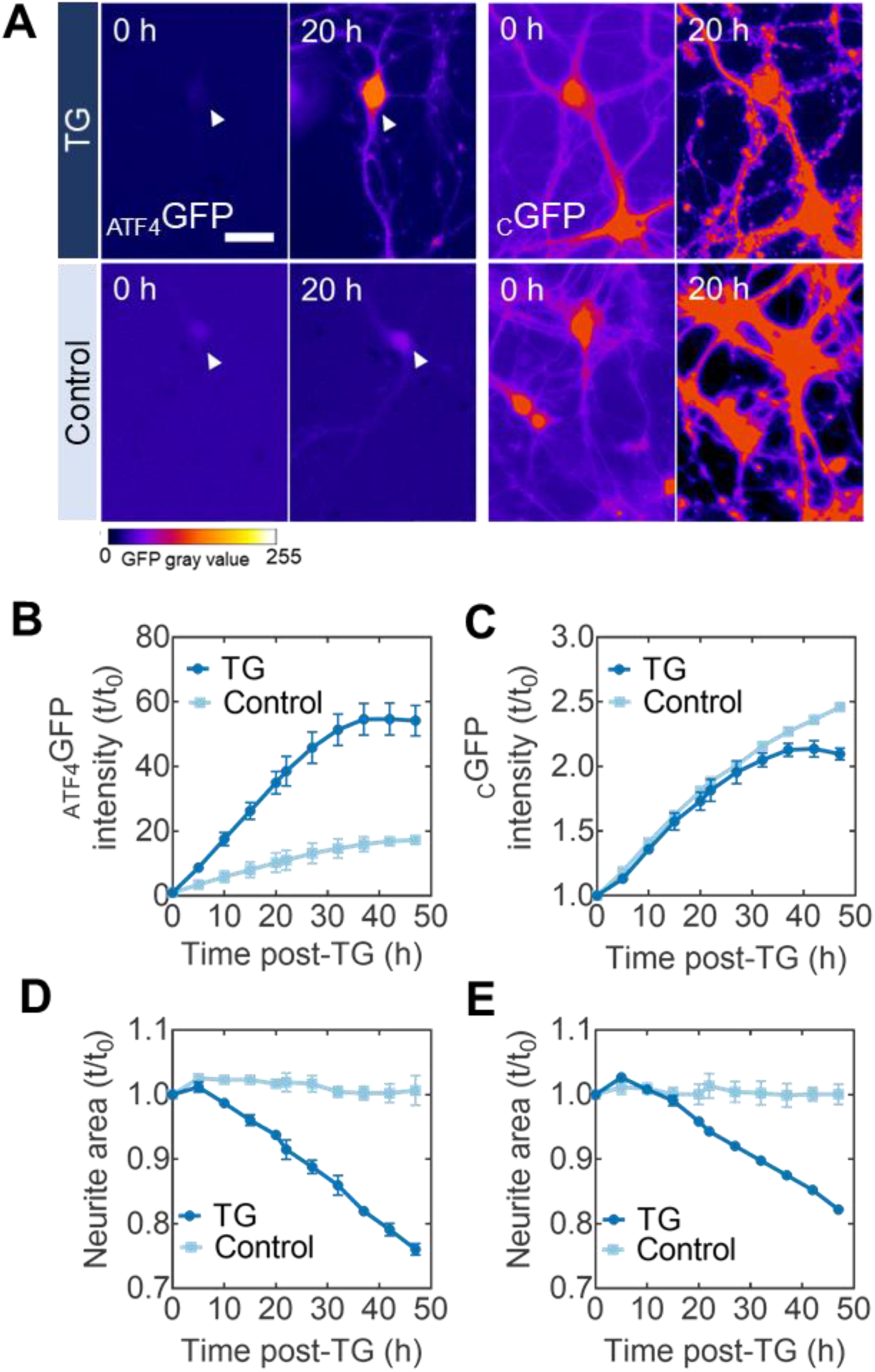
Kinetic monitoring of ER stress in live neurons. (**A**) Representative images of primary neurons transduced with _ATF4_GFP (left) or _C_GFP (right) at 0 h and 20 h post-exposure to either the ER stress inducer TG (top) or DMSO control (bottom). Heatmap false coloring is used to represent GFP gray values for better visualization of fluorescent expression. Scale bar = 50 µm. (**B**) Kinetic plots for the mean _ATF4_GFP integrated intensity induction levels after TG (circles) or baseline levels with control (DMSO) treatment (squares) throughout the time-lapse imaging session. (**C**) Kinetic plots for the mean _C_GFP integrated intensity expression levels with TG (circles) or control (DMSO) treatments (squares). (**D-E**) Quantification of neurite area (neurite area = total area – cell body area) following TG-induced stress in either _ATF4_GFP or _C_GFP cultures. TG used at a final concentration of 100 nM. Error bars report SEM from biological triplicates. All kinetic plots are normalized to the initial reading t_0_.

The temporal profile of _ATF4_GFP expression over 48 h revealed a three-fold induction in _ATF4_GFP integrated intensity (the product of mean GFP intensity and area) compared to DMSO-treated, control cultures (Figure 2B). To ensure this rate of increase was a stress-dependent response, control sister cultures were transduced with _C_GFP and subjected to the same treatments. Neurons with _C_GFP showed similar expression rates regardless of treatment conditions over the 48-h period of experimentation (Figure 2C), consistent with the lack of a stress-responsive element in the cGFP construct. Concomitant with the induction of _ATF4_GFP, we found a reduction (∼20% total after 48 h) in neurite area following the TG treatment (Figure 2D, 2E), resembling neurodegeneration of the primary neurons. A morphometric analysis of β-tubulin III-stained processes confirmed the neurodegeneration-like effect of TG-induced ER stress across neurite metrics such as branching and terminal endpoints (Figure S2).

### Neuronal stress response to proteasome inactivation

Together with the autophagy-lysosomal pathway, the ubiquitin-proteasome system serves as a critical node of the proteostasis network by regulating protein degradation. Deficits in protein turnover due to proteasome impairment can elicit a condition of cellular stress and triggers the mobilization of the UPR signaling program. Importantly, proteasome impairment by cell-autonomous factors (e.g. protein oligomers) or disease co-factors (e.g. aging) has been implicated in the pathophysiology of several neurodegenerative disorders (Deger et al., 2015). Prior work has shown that proteasome inactivation by MG132, a pharmacological inhibitor of the proteasome, activates PERK in primary neurons and Neuro-2a cells consistent with ER stress and UPR activation (Zhang et al., 2010). We combined live-cell imaging with global inactivation of the proteasome by MG132 to gain a deeper insight into the relationship between the temporal characteristics of the neuronal stress response and cell death. Additionally, employing a pharmacological inhibitor of proteasome function allowed us to probe neuronal stress in a condition of proteostasis impairment not directly related to ER-targeted perturbations such as TG.

In primary cortical neurons, phenotypic characterization via immunocytochemistry following 24-h exposure to MG132 confirmed a dose-dependent decline in cell viability as well as an increase in ubiquitin positive staining (Figure S3A, S3B). We additionally quantified the effect of proteasome inactivation on tau-1, a neuronal protein client of the proteasome (David et al., 2002), and found a dose-dependent shift of expression patterns from smooth to punctate (Figure S3C), indicating disruption in processing of tau-1. Based on these results, we selected 100 nM for time-lapse fluorescence imaging experiments. Upon treatment, we found that exposure to MG132 caused a robust increase in _ATF4_GFP in neurons with maximum levels observed at 26-h post-treatment followed by a signal decline consistent with cell loss (Figure 3A, 3B).

**Figure 3:**
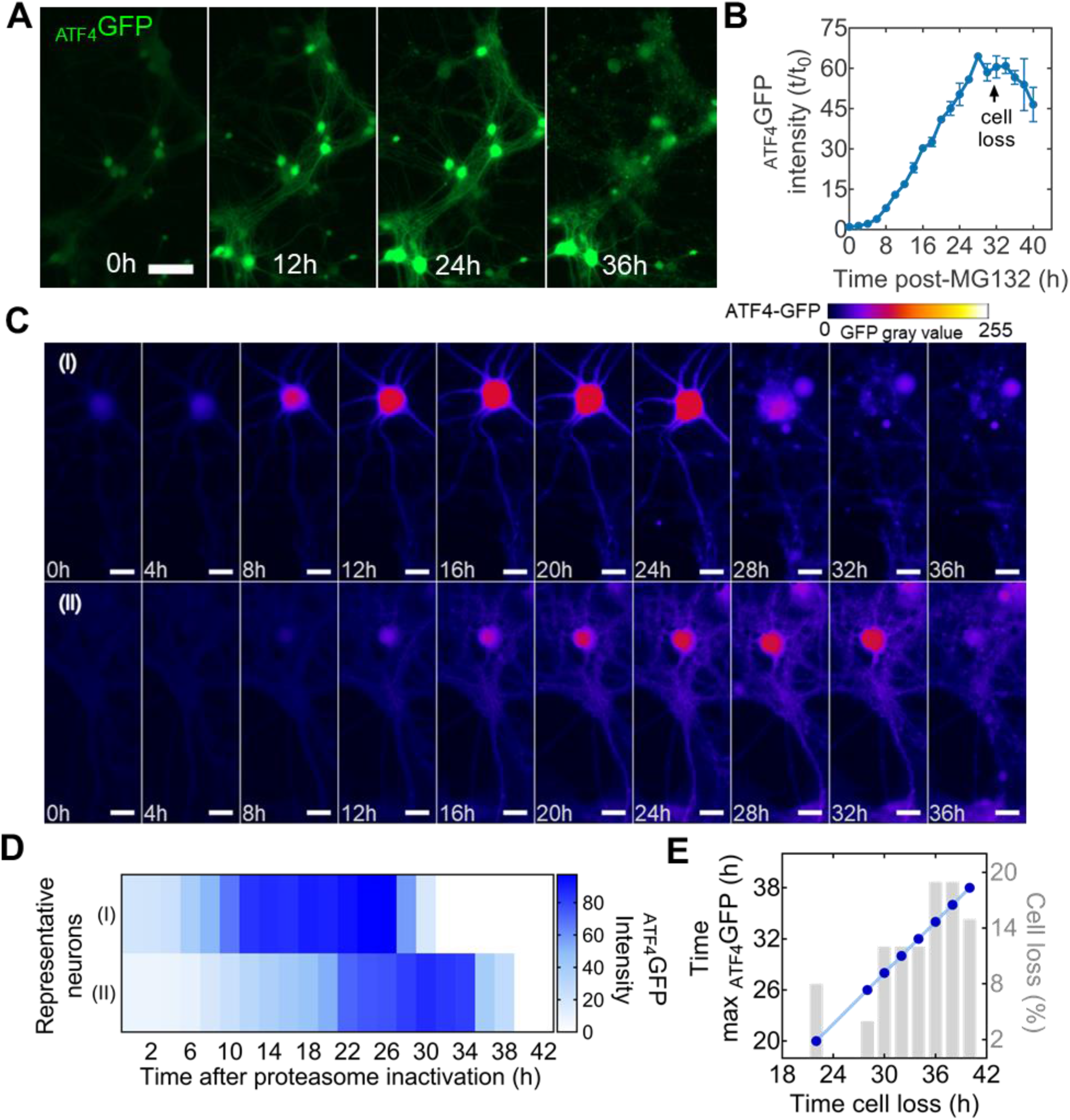
Population and individual kinetic detection of cellular stress following proteasome inactivation. (**A**) Representative time-lapse microscopy images of _ATF4_GFP induction in neurons after treatment with 10 nM of the proteasome inhibitor MG132. Scale bar = 100 µm (**B**) Kinetic plot for the mean _ATF4_GFP integrated intensity over the course of 40-h following MG132 treatment. Arrow indicate inflection point associated with cell loss. Error bars report SEM from biological triplicates. (**C**) Individual neurons from the same population show different rates of _ATF4_GFP induction and time of cell death (loss of membrane integrity). Heatmap false coloring is used to represent GFP gray values for better visualization of fluorescent expression. Scale bar = 25 µm. (**D**) Intensity visualization plot for each neuron in (C) describing intensity of _ATF4_GFP expression at each time point. (**E**) Correlation plot between the time at which neurons showed maximum _ATF4_GFP expression (left y-axis) and the time of their eventual loss (x-axis). Percentage of total cell death for the corresponding time point (right y-axis).

Through conventional biochemical and molecular biology assays, UPR induction in neurons has been primarily described at a population level. However, a fundamental concept in cell biology is that individual members of a cell population can exhibit distinct biological responses (Niepel et al., 2009). We leveraged the capability of probing UPR-related stress responses by live-cell microscopy to visualize individual neuronal responses to proteasome inactivation by MG132. While the typical response, where an increase in _ATF4_GFP was followed by loss of membrane integrity (cell loss), was consistent across the analyzed cells from the same population, the time of induction and cell death differed between individual neurons (Figure 3C). A fundamental notion of the UPR is that in conditions of unresolved stress, this homeostatic mechanism switches into a terminal cell death signaling program. Confirming this, we found that delayed stress induction coincided with later time of cell loss (Figure 3D) with ∼54% of surveyed neurons dying between 34 and 40 h post-MG132 exposure (Figure 3E). Furthermore, timings of the maximum _ATF4_GFP induction and the subsequent cell loss determined in individual cells showed a strong positive correlation (Figure 3E). Our analysis supports the notion that the time for each neuron to reach maximum _ATF4_GFP expression can predict neuronal death, even if the timing of proteasome inactivation may vary among cells.

### Mutant α-synuclein overexpression elicits and influences neuronal stress response to proteostasis impairment

The alanine-to-threonine point mutation at the amino acid position 53 in α-synuclein (α-syn(A53T)) is implicated in familial forms of Parkinson’s Disease (PD) (Polymeropoulos et al., 1997). Biochemical studies have shown that the point mutation promotes aggregation of α-syn protein (Conway et al., 1998). The presence of α-syn(A53T) is known to disrupt proteostasis either by directly engaging the UPR and ER stress signaling, hindering protein degradation pathways, or by indirectly overworking protein chaperone machinery. As a disease model, overexpression of α-syn(A53T) is known to cause α-syn-rich protein inclusions and neurodegeneration (Ip et al., 2017; Li et al., 2013).

Overexpression of α-syn(A53T) in neurons impacts proteostasis by increasing the burden of protein synthesis, particularly the synthesis of an intrinsically disordered protein. To visualize neuronal stress responses to such proteostasis dysfunction, we overexpressed α-syn(A53T) and monitored _ATF4_GFP under different experimental conditions. We first asked whether α-syn(A53T) overexpression alone was sufficient to elicit detectable levels of _ATF4_GFP in neurons. We observed that at 10 d after lentiviral transduction, _ATF4_GFP levels were significantly higher in neurons overexpressing α-syn(A53T) than in negative controls (Mann Whitney test; *p*=0.0002; *n*=8 biological replicates; Figure 4A, 4B). Thus, similar to prior work demonstrating UPR induction in the *SNCA* triplication model of PD (Heman-Ackah et al., 2017), our results suggest UPR engagement in neurons overexpressing α-syn(A53T).

**Figure 4:**
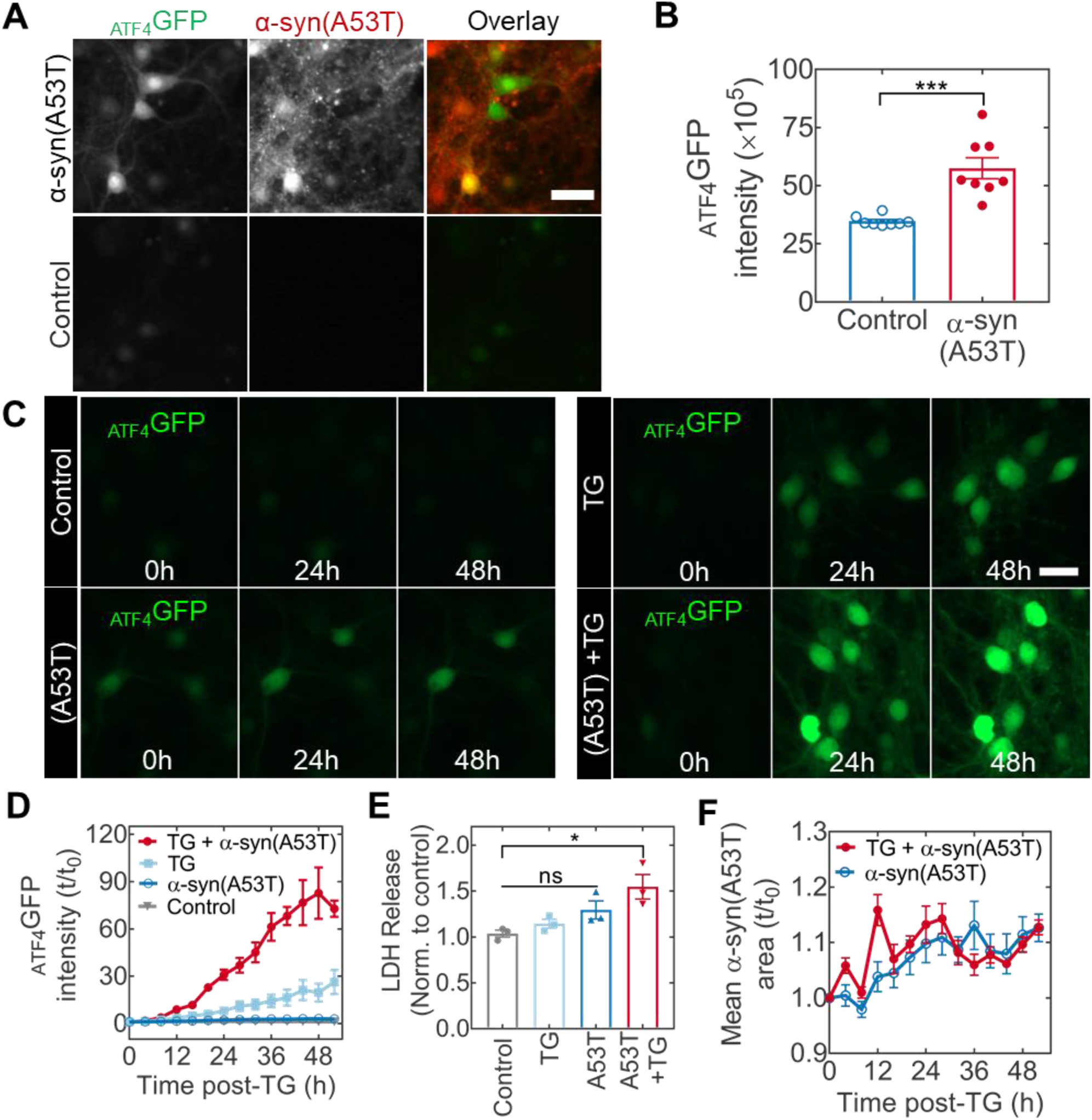
Mutant α-syn(A53T) overexpression elicits cellular stress and influences the neuronal response to ER stress. (**A**) Induction of cellular stress in primary neurons at 10 d after the co-transduction with _ATF4_GFP and mKate2-tagged α-syn(A53T) vectors. (**B**) Quantification of _ATF4_GFP integrated intensity in neurons with (red) or without (blue) α-syn(A53T) expression. \*\*\**p* = 0.0002 by two-tail Mann-Whitney test. (**C**) Representative time-lapse microscopy images from neurons expressing _ATF4_GFP and left untreated (top left) or subjected to α-syn(A53T) overexpression (bottom left), TG stress (top right), or a combination of α-syn(A53T) and TG (bottom right). Scale bar = 50 µm. (**D**) Corresponding kinetic plots for the mean _ATF4_GFP integrated intensity for all groups. (**E**) Assessment of cellular health via LDH quantification in cell media. Absorbance values are normalized to an untreated control. Multiple comparisons by Kruskal-Wallis test (untreated vs A53T+TG \**p* = 0.0395; rest not significant. (**F**) Quantification of mean α-syn(A53T) area (red fluorescence positive puncta) with (closed circles) or without (open circles) TG stress.

We then asked whether α-syn(A53T), an aggregation-prone, non-ER client protein, could influence neuronal susceptibility to ER stress. In addition to prominent protein misfolding, ER stress is considered a disease co-factor in many neurodegenerative conditions. Therefore, the temporal response of neurons subjected to such level of combined proteostasis dysfunction remains a key question in the understanding of how α-syn(A53T) contributes to neurodegenerative outcomes. Following a 50-h time-lapse study, representative microscopy images show induction of UPR captured by _ATF4_GFP expression in α-syn(A53T), TG, and TG with α-syn(A53T) treated neurons (Figure 4C). The additional induction of _ATF4_GFP by α-syn(A53T) was more than two-fold compared to the TG treatment alone (Figure 4D). Notably, while _ATF4_GFP induction in response to α-syn(A53T) by itself was evident (Figure S4, insert), the induction level remained well below the effect of TG. To further assess the impact of α-syn(A53T) on TG-induced stress, we assayed for LDH release into the culture media as an indicator of cellular health. We found that the presence of α-syn(A53T) sensitized neurons to LDH release following treatment with TG (Figure 4E). Altogether, we conclude that overexpression of α-syn(A53T) can predispose neurons to increased vulnerability in conditions of ER stress.

To further look into the interplay between α-syn(A53T) and ER stress, we hypothesized that ER stress could lead to increased intracellular aggregation of α-syn(A53T). In yeast, ER stress caused by reducing conditions (dithiothreitol) or glycosylation inhibition (tunicamycin) is sufficient to induce the aggregation of cytosolic, non-ER targeted, proteins (Hamdan et al., 2017). In the presence of tunicamycin, α-syn shows enhanced aggregation propensity in dopaminergic neurons (Cóppola-Segovia et al., 2017). To evaluate this notion but with TG-induced ER stress, we quantified fluorescence coverage area by α-syn(A53T) following the TG treatment by tracking the mKate2 fluorescence tag. Contrary to other inducers of ER stress, our results show similar fluorescence area over time in the presence or absence of TG (Figure 4F), suggesting that the ER stress does not necessarily promote aggregation of α-syn(A53T).

## DISCUSSION

In the current study, we developed and characterized stress-dependent expression vectors to monitor proteostasis dysfunction in neurons. We first showed how an AAV-based reporter constructs can harness the selective translation of *ATF4* mRNA as part of the induction of the UPR to yield a quantifiable readout suitable for non-cell destructive monitoring of cellular stress due to calcium dysregulation in the ER by TG. We extended our characterization studies to non-ER targeted perturbations, namely proteasome inhibition by MG132, and present data that describes the temporal profile of stress progression and cell loss in neurons experiencing global inactivation of the proteasome machinery. Finally, we adopted the overexpression of α-syn(A53T) in neurons, an established *in vitro* model of PD, and describe the impact of α-syn(A53T) on eliciting cellular stress and influencing neuronal susceptibility to ER stress.

Proteostasis dysfunction is an important manifestation in all major neurodegenerative diseases. However, while analysis of proteostasis dysfunction via fluorescent reporters of UPR activation has been previously reported (Lajoie et al., 2014), these molecular tools have not been incorporated into neurobiologically relevant contexts. Herein, we reported our initial efforts towards adapting such technologies for use in neurobiological research. Using mouse primary neurons as a cellular system, our results demonstrated the induction characteristics of _ATF4_GFP expression as a surrogate for ER stress and UPR activation in a cell-based assay suitable for high-throughput screening studies. Cell-based assays amenable to non-destructive longitudinal monitoring, and with the dynamic range to either capture acute (e.g. thapsigargin) or chronic (e.g. α-synuclein) stress responses, can have an impact on neuroscience discovery research and beyond. For example, such assays can facilitate a more refined understanding of the mechanisms of action as well as time-cellular fate relationships in the search for pharmacological modulators of the UPR. Besides neurodegeneration, UPR activation, particularly PERK signaling, is central to the development of human diseases like pancreatic β-cell dysfunction in diabetes, inflammation, atherosclerosis, and diabetes (Wang and Kaufman, 2012). Therefore, a wide range of applications exists for which a cell-based assay for stress detection compatible with primary cell sources and temporal monitoring would be of great benefit.

The ubiquitin-proteasome system is one of the two identified protein degradation pathways in eukaryotic cells contributing to the efficient degradation and turnover of proteins. Pharmacological blockade of the proteasome is an established perturbation to proteostasis leading to cellular stress and eventually cell death. In HeLa cells, proteasome inhibition with MG132 results in stress granules in a response coordinated by the GCN2-eIF2α axis of the ISR (Mazroui et al., 2007). In contrast, prior work in primary neurons and a neuronal cell line identified the activation of PERK as an event in the cellular response to different types of proteasome inhibitors (Zhang et al., 2010). In addition to supporting the notion of UPR engagement, our work provides kinetic data to describe the progression of neuronal stress following proteasome inhibition in neurons.

Additionally, consistent with previous findings in non-neuronal systems, our results indicate a clear relationship between the temporal progression of stress induction and cell death. Notably, visualizing asynchronous cell death within a cell population, a process in which only 5-10% of cells will execute apoptosis at any given moment, represents a technical challenge (Mills et al., 1998). By examining neuronal stress using live-cell imaging, we distinguished neuronal death asynchronism and correlated the time of cell loss with maximum stress levels. The relationship between stress progression and induction of a specific mechanism of cell death remains to be explored. Consistent with our measurement of an ATF4-based reporter signal, the most likely cell death mechanism following sustained UPR activation is mediated by CHOP, an ATF4-target gene that orchestrates apoptosis (Galehdar et al., 2010). However, given the acute nature of MG132 treatment, we cannot exclude the potential contribution of other factors such as oxidative stress in the cytotoxicity that we observed.

The presence and accumulation of misfolded species of α-synuclein are recognized as important contributors to neurodegeneration in Parkinson’s Disease (PD) as well as neurodegenerative conditions collectively known as α-synucleinopathies. In addition to PD, major α-synucleinopathies include Dementia with Lewy Bodies (DLB) and Multiple Systems Atrophy (MSA). In each condition, it is believed that α-syn pathogenicity stems from conformational changes that lead to fibrillar cellular inclusions referred to as Lewy pathology of which α-syn is the main component (Adamowicz et al., 2017; Spillantini et al., 1998). When in excess, such as a result of duplication or triplication of the gene encoding α-syn (*SNCA*), wild-type α-syn can trigger early-onset PD (Goedert et al., 2013; Siddiqui et al., 2016). Additionally, human genetics have identified six mutations in *SNCA* to be linked to familial forms of PD. In particular, the A53T point mutation in α-syn is linked to early-onset PD in humans (Polymeropoulos et al., 1997). At a molecular level, mutant α-syn(A53T) displays faster fibrilization and aggregation behavior compared to wild-type α-syn (Conway et al., 1998).

We overexpressed mutant α-syn(A53T) in wild-type background of primary neurons to determine whether our _ATF4_GFP reporter could capture stress responses arising from such manipulation to neuronal proteostasis. While the overexpression model of α-syn(A53T) does not yield insoluble inclusions *in vitro*, multiple lines of evidence suggest that α-syn oligomers can elicit toxicity without the need of forming high molecular weight aggregates (Prots et al., 2018; Winner et al., 2011). Our studies found that the overexpression of α-syn(A53T) alone could produce detectable levels of stress 10-d post-transduction. How this level of stress response relates with known manifestations of α-syn(A53T) such as mitochondrial dysfunction (Ordonez et al., 2018) or axonal degeneration (Prots et al., 2018) warrants further investigation.

The study of the interaction between α-syn and other disease co-factors is an active area of research. Given the intricacies of the pathophysiology of conditions such as PD, the most likely description of disease pathophysiology includes the combination of aggregation-prone proteins such as α-syn and co-factors, including aging, ER stress, and neuroinflammation. It has been hypothesized that proteinaceous inclusions in neurodegenerative diseases could be a consequence of an overall decline in proteostasis due to aging (Klaips et al., 2018). Proteostasis function can also be affected by other factors such as neuroinflammation via increased ER stress (Pintado et al., 2017). We examined whether the presence of α-syn(A53T) influenced the degree of the neuronal response to ER stress. Whether the presence of α-syn(A53T) lowers the threshold or simply amplifies UPR signaling remains to be fully elucidated. However, our data indicate that, with α-syn(A53T) overexpression, neurons elicited a more robust stress response to ER stress induced by thapsigargin. Furthermore, our results elucidated increased vulnerability where cellular health was compromised due to the combined effect of α-syn(A53T) and thapsigargin-induced ER stress in neurons. This observation is consistent with previous studies indicating that, in the context of PD, a secondary insult (e.g., environmental factor) is needed to induce cell death (Chang et al., 2016).

Future iterations of the approach presented herein should focus on certain opportunities for extended functionality. For instance, to study the temporal induction of organelle-specific UPR signaling such as mitochondrial UPR (reviewed by Shpilka and Haynes, 2018), this ATF4-based reporter could be modified with organelle-specific targeting sequences. Similarly, previous work has described compartment-specific (cell body vs. processes) stress responses directed by the local translation of ATF4 in axons (Baleriola et al., 2014). A construct design incorporating cell compartment-specific localization signals would allow for such subcellular analyses. Finally, by introducing cell-type specific promoters, stress-responsive AAV constructs could enable a direct assessment of how different neuronal subtypes respond to proteostasis impairment, further illuminating specific neuronal vulnerabilities in disease. Hence, the reporter construct presented in this work can enable stress-dependent labeling of distinct neuronal populations in the CNS following chemical or genetic-based stressors *in vivo*.

## CONCLUSIONS

In conclusion, this work presents the development and characterization of stress-dependent expression vectors that enable the longitudinal study of neuronal responses to ER stress and proteostasis dysfunction in live neurons (individual or population). Using this reporter assay we show that during unresolved stress, neuronal death is preceded by a relative maximum induction of the UPR. Additionally, we describe how the Parkinson’s-related mutant protein α-syn(A53T) influences how neurons respond to ER stress. Collectively, this work bridges the gap between advances in the live-cell analysis of cellular responses to proteostasis dysfunction and their application in neurobiology. By adopting an experimental approach like the one presented herein, we hope this work will stimulate the study of mechanisms of neurodegeneration associated with the induction of neuronal stress responses.

## ACKNOWLEDGEMENTS

This study was supported by an NIH-NINDS Predoctoral Fellowship T32 NS007480-18 (AJS-L), the Emory University Research Committee (REG and CAG), NIH R21 NS112740-01 (REG), and S10 OD021773 (KB).

---

**Figure S1:**
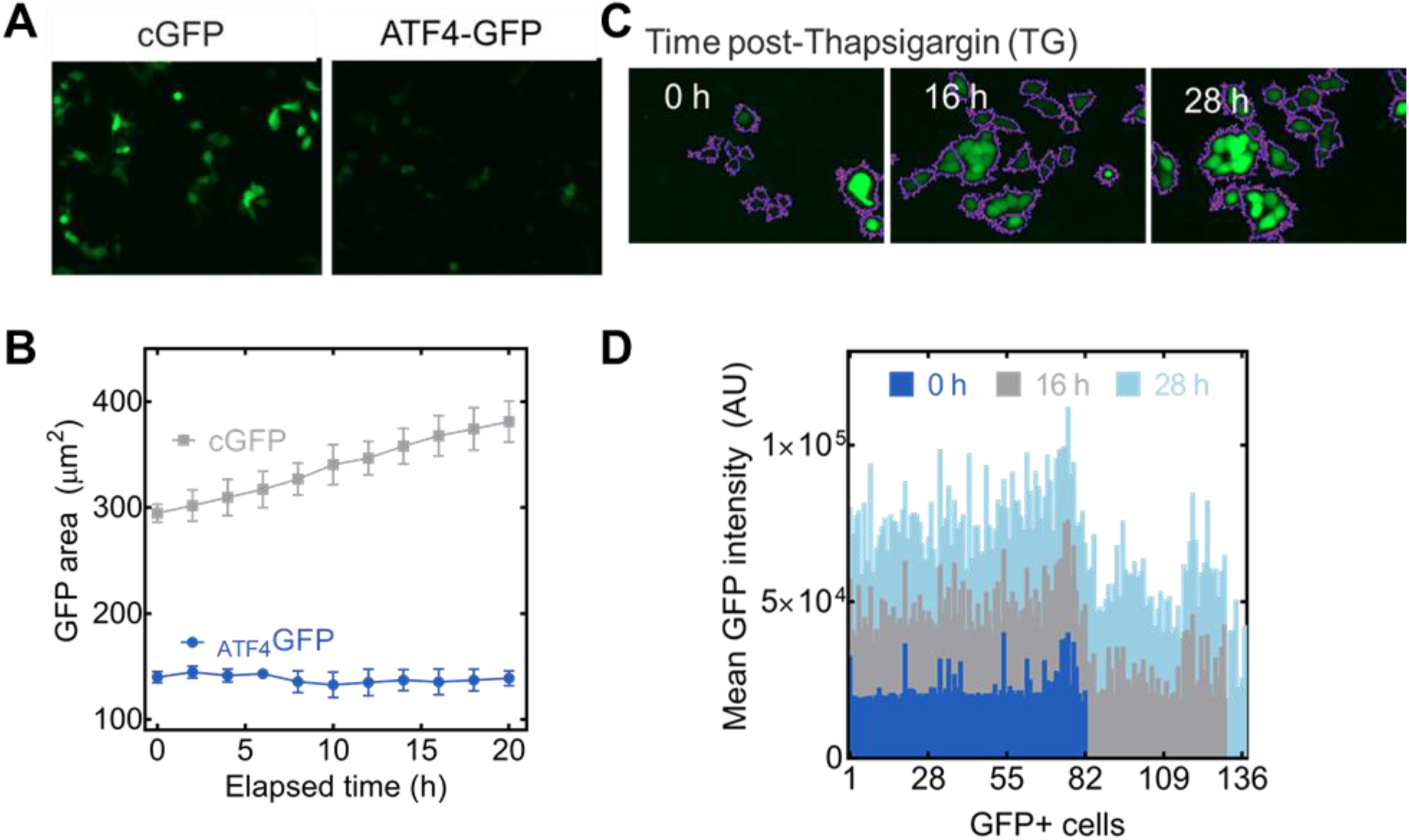
(**a**) Suppression of reporter expression evaluated 48 h post-transfection. (**b**) Kinetic monitoring of expression stability over 20 h. (**c**) and (**d**) Induction of reporter expression following treatment with Thapsigargin (TG).

**Figure S2:**
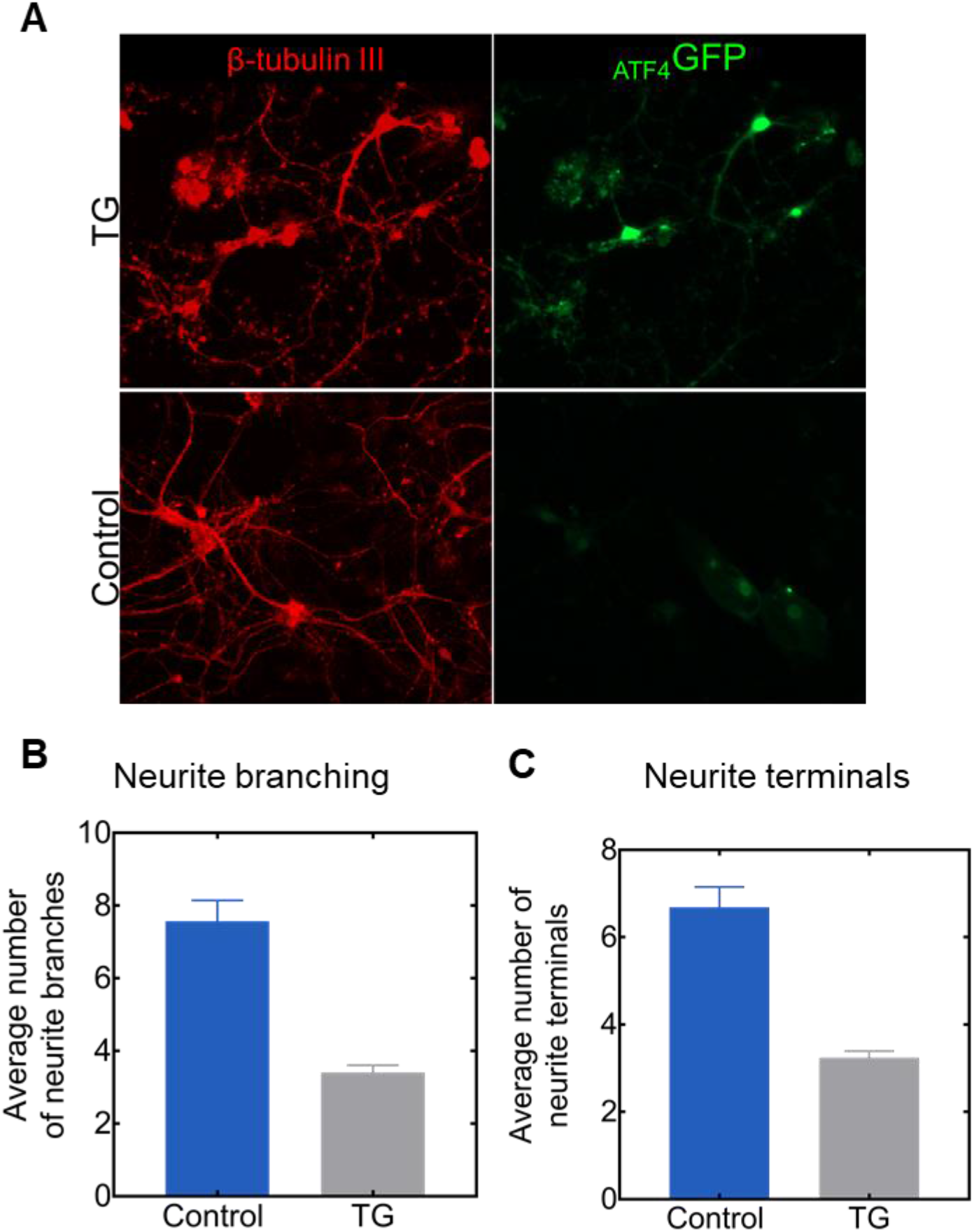
Endpoint morphometric analysis of primary neurons following 48 h exposure to the ER stressor thapsigargin (TG). (A) Cortical neurons immunostained for β-tubulin III (TUJ1) to label projections. (B) and (C) Quantification of neurite branching and terminals as measured by the CellProfiler software. A custom-made algorithm used a morphological marker (e.g., TUJ1) to threshold neurons in each field. The next step identified soma (cell bodies) as primary objects based on size. Neurites were then identified as secondary objects (nested within the primary objects step) and subjected to a smoothing filter. The resulting image was then ‘skeletonized’ using the MorphNeurites processing step. As an output, this process generated data sets for neurite length, branches, and end points (terminals) which facilitated quantifying the effect of TG-induced ER stress in neurons.

**Figure S3:**
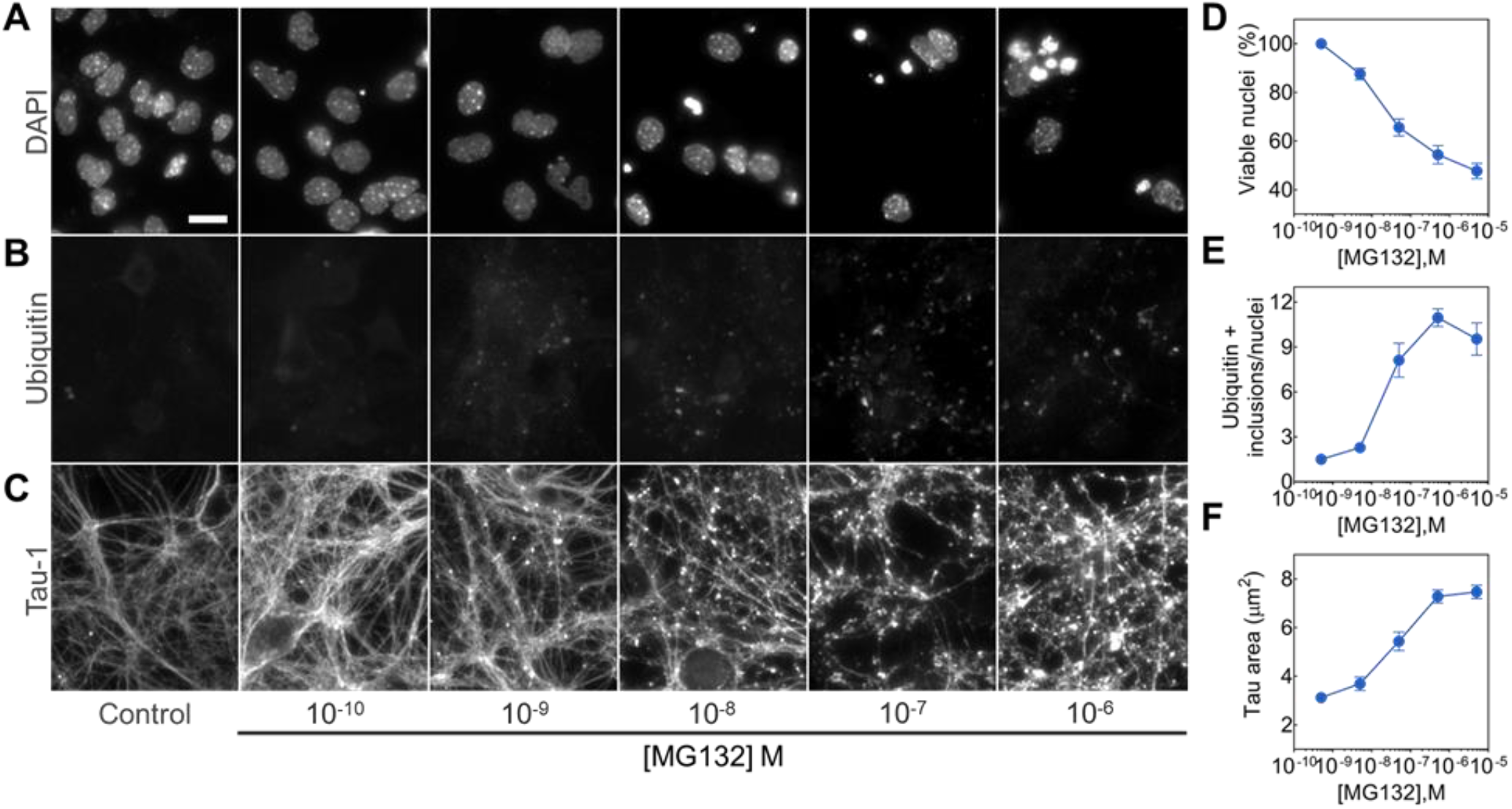
Phenotypic characterization of the effects of MG132 proteasome inhibition in primary neurons. (**A**) Dose-dependent decline in viable nuclei as determined by image analysis of DAPI staining. (**B**) Accumulation of ubiquitin positive inclusions. (**C**) Disrupted expression patter of the axonal protein tau following MG132 treatment. (**D-F**) Corresponding quantification based on n=4 biological replicates for each concentration.

**Figure S4:**
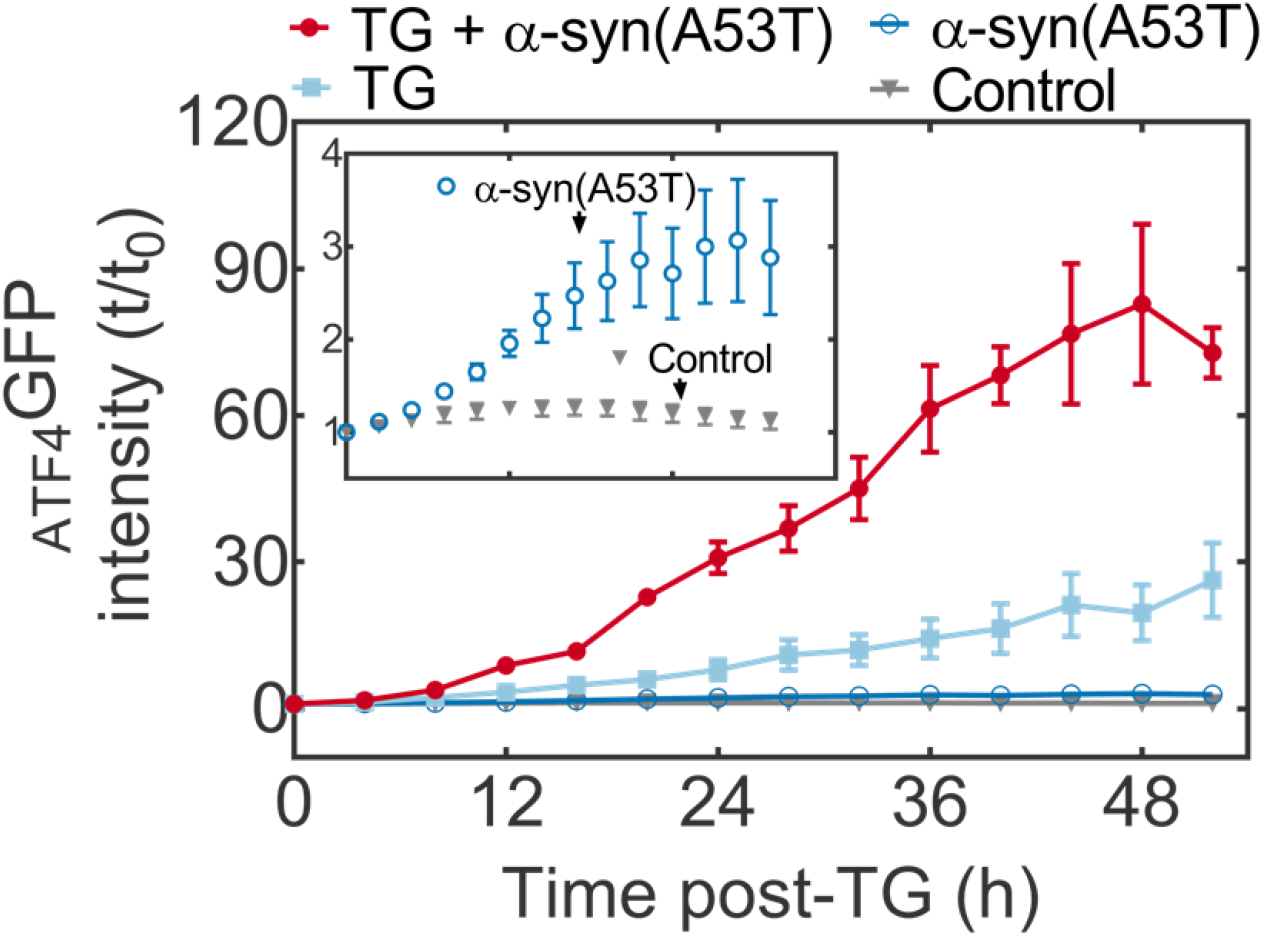
Corresponding kinetic plots for the mean _ATF4_GFP integrated intensity from neurons expressing _ATF4_GFP and left untreated or subjected to α-syn(A53T) overexpression (insert) and neurons expressing _ATF4_GFP and exposed to TG stress or a combination of α-syn(A53T) and TG (main plot).

